# Nutrient Availability Regulates an Exploration-Elaboration Trade-off in Fungal Rhizomorph Networks

**DOI:** 10.64898/2026.08.23.746505

**Authors:** Sarah C. Naeher, Markus J. Buehler

## Abstract

Rhizomorphs are specialised, root-like fungal structures whose hierarchical organisation may offer a route to reinforcing mycelium-based materials, yet the environmental regulation of their network formation remains poorly understood. Here, we develop an image-based framework to characterise the longitudinal growth and organisation of *Armillaria gallica* rhizomorphs under varying nutrient availability and light exposure. Time-lapse imaging was combined with image segmentation, skeleton-based network analysis, optical measurements and Gompertz growth modelling. Nutrient availability produced a distinctly non-monotonic response. Moderate nutrient limitation (0.5*×* standard concentration) favoured rapid and coherent exploratory growth, whereas intermediate enrichment (1.5*×*) produced the greatest eventual network extent, reaching approximately 975 mm total strand length; network extent and radial expansion differed significantly across nutrient levels (*p*_adj_ = 0.0012). Further enrichment maintained substantial fungal coverage without additional network elaboration, consistent with a shift from long-range exploration towards more locally consolidating growth. By contrast, exclusion of ambient light produced no significant differences after multiple-testing correction. These results reveal a resource-dependent trade-off between exploration and network elaboration, demonstrate that fungal coverage and organised network formation are distinct outcomes, and provide a quantitative basis for controlling self-organised biological architectures for bio-derived material design.

## 1. Introduction

Fungal networks provide a natural model for how living systems construct spatially distributed structures that adapt their morphology to local environmental and resource conditions while supporting exploration, transport, and survival. Understanding the principles governing the emergence and organisation of such networks therefore connects questions in organismal biology and network dynamics with opportunities in bio-inspired engineering and living materials [8]. Fungal mycelia form adaptive transport networks whose architecture reorganises in response to resource availability and environmental conditions, coupling growth and network remodelling with the redistribution of matter across space [3, 14]. More broadly, such self-organising biological architectures illustrate how functional material organisation can emerge through local environmental feedback, connecting fungal biology with network science and concepts of biological material intelligence [23].

Construction and packaging materials contribute substantially to global CO_2_ emissions, motivating the development of sustainable alternatives with lower environmental impact [1]. Among these, mycelium-based composites (MBCs), produced by growing fungal mycelium through lignocellulosic substrates such as agricultural residues, have attracted increasing interest because of their low density, biodegradability, thermal insulation, hydrophobicity, and fire-resistant properties [12]. Their broader application, however, remains constrained by comparatively weak and variable mechanical performance, particularly for structural or load-bearing uses [1]. Recent fabrication approaches, including additive manufacturing of biopolymer composites followed by fungal inoculation, have expanded the geometric design space and enabled greater control over material architecture [32]. Nevertheless, improving the mechanical performance of fungal materials will also require a better understanding of how fungi form and organise mechanically robust structures. Biomateriomics provides a framework for this effort, treating processing–structure–property relationships in biological materials as quantifiable and ultimately designable [8, 11, 15, 24, 31, 39]; the present work addresses the first of these links, from cultivation environment to network organisation.

Rhizomorphs represent a potentially valuable biological system in this context. Fungal hyphae can differentiate into mycelial strands, cords, and rhizomorphs, with rhizomorphs forming as highly organised, root-like organs associated with nutrient exploration and long-distance resource translocation [2, 6, 18]. Network-level analysis of naturally occurring *Armillaria* rhizomorph systems has further shown that connectivity and redundancy can facilitate resource redistribution and confer robustness to local network disruption [21]. In contrast to relatively simple mycelial cords, in which constituent hyphae remain more loosely associated, rhizomorphs consist of large numbers of interlaced and adherent hyphae organised into differentiated tissues [36–38]. In *Armillaria gallica*, these tissues are arranged radially into peripheral hyphae, a cortical layer, longitudinally oriented medullary tissues, and a central cavity [38]. This differentiated architecture supports specialised functions including long-distance transport of water and nutrients and internal gas conduction during growth through poorly aerated substrates [2, 10, 17, 25]. Mature *Armillaria* rhizomorphs may additionally develop a melanised, calcium-containing outer layer that provides mechanical and chemical protection [28]. Their capacity for invasive growth is further supported by substantial turgor-driven force generation, allowing rhizomorphs to penetrate mechanically resistant substrates [37, 38]. These structural and functional characteristics suggest that rhizomorphs may provide greater mechanical robustness than undifferentiated mycelial networks and therefore warrant investigation as potential reinforcing structures in fungal materials.

Rhizomorph initiation and extension are influenced by a broad range of interacting physical, chemical, and atmospheric conditions. Temperature is an important determinant of development, with several studies reporting favourable growth within an approximate range of 20–26 °C and reduced development at higher temperatures [26, 27, 30]. Water availability can influence extension both directly and through interactions with temperature and gas exchange [27, 33], while ambient pH affects rhizomorph differentiation over a somewhat narrower range than that supporting general mycelial growth [34]. Oxygen availability and diffusion are important for sustained extension, whereas elevated oxygen concentrations at the rhizomorph surface can inhibit apical growth; CO_2_ appears to exert a comparatively limited effect over moderate concentrations [19, 29, 33]. Mechanical properties of the substrate can also influence development, with increased gel resistance reported to stimulate rhizomorph extension [37]. These findings illustrate the environmental sensitivity of rhizomorph development, but also demonstrate that the effects of individual factors may be difficult to isolate because environmental controls frequently interact.

Among these factors, nutrient availability is particularly closely linked to the ecological function of rhizomorphs. Rhizomorphs contribute to fungal foraging by enabling growth away from established nutrient sources while translocating resources towards advancing growth fronts [2, 6, 13, 17, 20]. Their initiation and development are sensitive to substrate nutrient status and, in particular, to the balance between carbon and nitrogen. In *Armillaria mellea*, intermediate carbon-to-nitrogen ratios have been reported to favour rhizomorph formation, whereas either excess or deficiency of these resources can inhibit development [34]. A minimum substrate nutrient status is also required for rhizomorph initiation, and establfsupplished rhizomorphs may suppress the formation of nearby independent initials through local nutrient depletion [16]. These observations suggest that nutrient availability may influence not only whether rhizomorphs form, but also the subsequent balance between rapid exploratory growth and the development of an extensive, differentiated network. However, the effect of overall nutrient density on the temporal dynamics, spatial expansion, and network architecture of rhizomorph growth remains insufficiently quantified.

Light represents a second potentially important environmental signal. Previous studies have suggested that rhizomorph formation is promoted under dark conditions [37], while illumination has been reported to reduce elongation in *Armillaria mellea* [9]. These observations indicate that light may act as an inhibitory stimulus for rhizomorphic development. However, the magnitude and nature of this response remain uncertain, particularly with respect to network-scale features such as colonisation kinetics, branching topology and pigmentation. Earlier studies have predominantly focused on initiation or elongation rather than longitudinal changes in the architecture of the developing network. More generally, quantitative comparison of rhizomorph growth across environmental conditions has been limited by the absence of scalable methods capable of tracking both colony-level expansion and network morphology over extended periods.

Image-based phenotyping provides a route to address this limitation. Recent advances in foundation models enable prompt-driven segmentation of complex images without task-specific model training [7], while multimodal AI increasingly provides mechanisms for extracting and reasoning over structural information contained in scientific images [5]. Applied to longitudinal imaging of fungal colonies, these approaches can transform visual observations into quantitative descriptions of spatial coverage, radial expansion and network architecture. Combining region-based segmentation with skeleton-based analysis further enables measurement of strand length, branching organisation and network connectivity, while optical measurements capture changes in colony appearance. Such complementary metrics are particularly important for rhizomorphs because development involves not only changes in the amount of fungal growth, but also changes in its spatial organisation, network topology and apparent differentiation. Beyond phenotyping, computational and generative AI approaches increasingly make it possible to translate structural information from biological systems into controllable material design spaces and ultimately into physical, fabricable geometries [22, 31].

Here, we investigate the longitudinal development of *Armillaria gallica* rhizomorphs under controlled laboratory conditions in two experiments examining nutrient density and light environment. Growth was monitored through repeated imaging and analysed using a computational workflow combining Segment Anything Model 3 (SAM 3) segmentation, region-based and skeleton-based morphological analysis, optical measurements, and Gompertz growth modelling. We asked how nutrient availability and the presence or absence of ambient light influence the timing, extent, spatial expansion, and organisation of rhizomorphic growth. In parallel, we evaluated the performance and limitations of the image-analysis framework across the different growth morphologies produced experimentally. By linking environmental conditions to longitudinal changes in rhizomorph development, this study provides a quantitative basis for understanding the regulation of organised fungal networks and for future investigation of their potential use as reinforcing structures in bio-derived materials.

## 2. Results and Discussion

The experimental and image-analysis framework was first assessed for its ability to capture reproducible rhizomorph growth dynamics. We then examined the effects of nutrient density and light environment. Within their respective experimental designs, nutrient density produced substantially broader changes in rhizomorph development, whereas exclusion of ambient light resulted in comparatively limited detectable differences.

### 2.1. Standard conditions produce reproducible growth dynamics

Under standard cultivation conditions, gel coverage followed a characteristic sigmoidal trajectory, consisting of an initial low-coverage phase until approximately 350 h, a period of rapid expansion between approximately 350 and 800 h, and a subsequent plateau. Gompertz models described the individual control trajectories well (*R*^2^ = 0.97–0.99), substantially outperforming linear fits (*R*^2^ = 0.55–0.77; see Supplementary Figure 1a). The timing of maximum growth was also consistent among replicates, with inflection points ranging from 475 to 528 h and a mean of approximately 499 h (see Supplementary Figure 1b). Between-replicate variability in gel coverage remained modest during both expansion and plateau phases, indicating that the standard cultivation regime produced highly reproducible colonisation dynamics.

**Figure 1.**
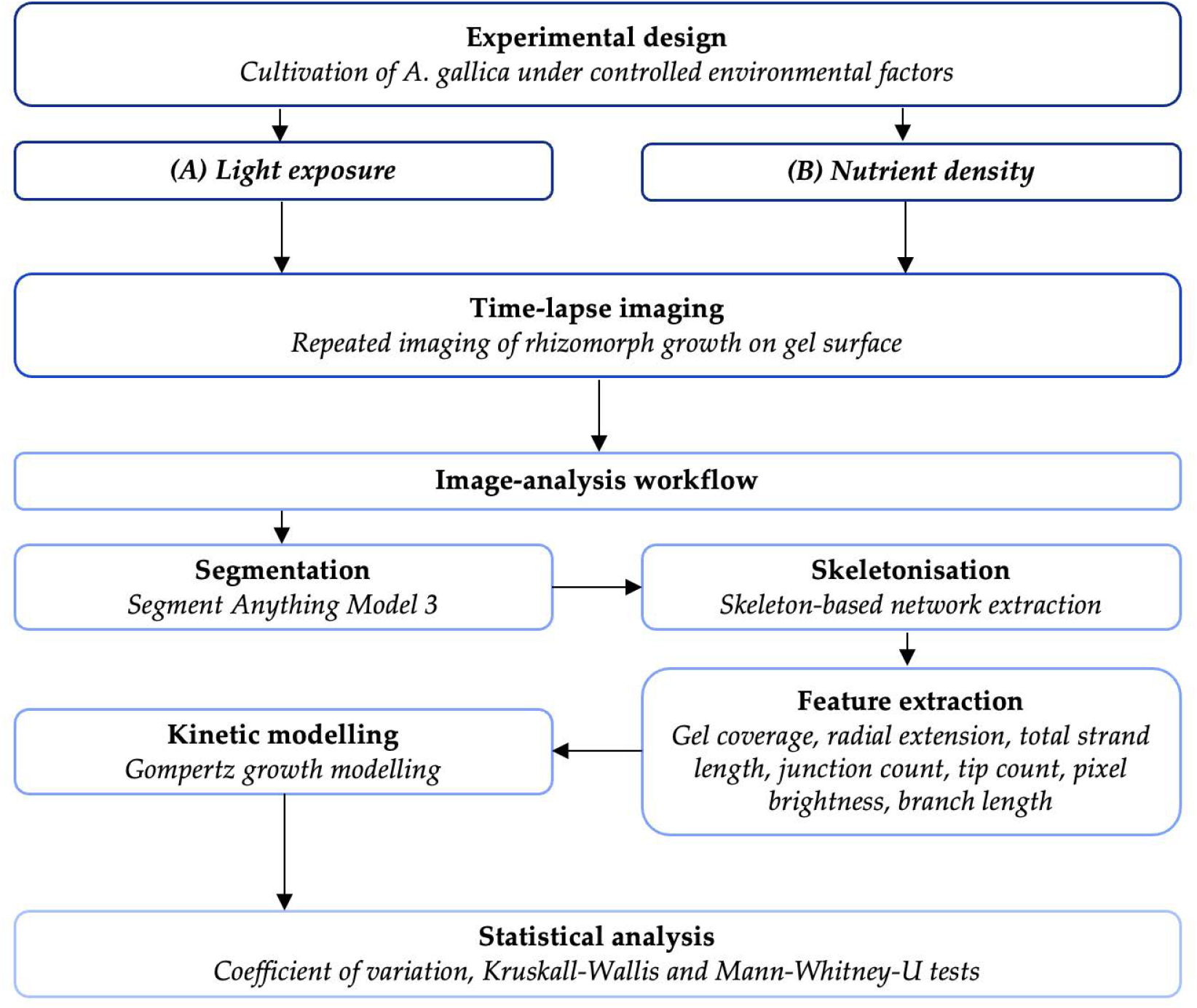
Schematic overview of the experimental and image-analysis workflow. Armillaria gallica was cultivated in two separate experiments examining light exposure and nutrient density, with rhizomorph development monitored by repeated time-lapse imaging. Images were segmented using Segment Anything Model 3 (SAM 3), followed by skeletonisation and extraction of morphological and optical features, including gel coverage, radial extension, total strand length, junction and tip counts, branch length, and pixel brightness. Gel-coverage trajectories were additionally characterised using Gompertz growth modelling, and differences among experimental conditions were evaluated using coefficients of variation and non-parametric statistical tests.

This reproducibility was also evident across morphological measurements in the light-exposure experiment (Figure 2). Most coefficients of variation were below 20%, with particularly low variability in mean radial extension and mean branch length. By contrast, several nutrient-density conditions exhibited substantially greater inter-replicate variability, particularly the most nutrient-limited condition and the intermediate enriched condition. The latter produced some of the most extensive networks observed in the study, but these responses were not equally expressed across all replicates. This variability is therefore relevant to the biological interpretation of the nutrient experiment rather than representing only measurement noise.

**Figure 2.**
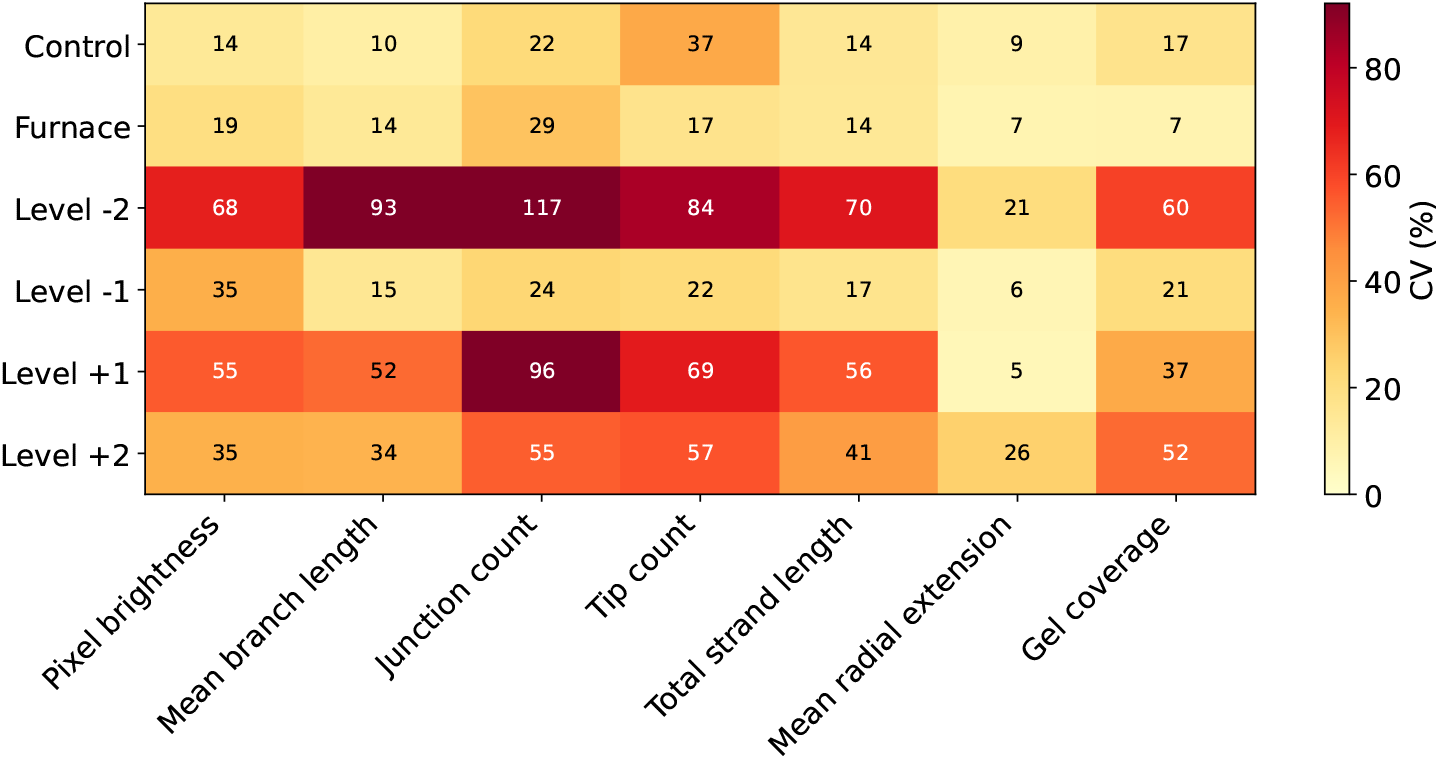
Reproducibility of rhizomorph growth across experimental conditions. Coefficients of variation (CV, %) were calculated across biological replicates for each morphological metric at the final shared timepoint; lower values indicate greater reproducibility. The ambient-light and dark-grown conditions showed consistently low variability, whereas several nutrient-density conditions were substantially more heterogeneous, indicating that nutrient availability affected not only mean growth behaviour but also its reproducibility across replicates.

The image-analysis workflow performed most reliably when fungal growth retained a clearly defined rhizomorphic morphology. Region-level segmentation showed good agreement with manually generated ground-truth masks under the standard light and dark conditions, whereas segmentation performance was more variable under the nutrient treatments. Importantly, area-based and length-based measurements were generally more robust than measures derived from exact skeleton topology. Junction and branch recovery became less reliable when growth was either diffuse or highly elaborate. Consequently, gel coverage, radial extension and total strand length provide the strongest quantitative basis for the comparisons below, while exact tip and junction counts are interpreted more cautiously.

### 2.2. Nutrient density non-monotonically regulates rhizomorph network formation

Nutrient availability substantially altered the timing, spatial extent, organisation and optical appearance of fungal growth. Six metrics differed significantly among the four nutrient levels after Benjamini–Hochberg correction: mean radial extension, total strand length, mean pixel brightness, tip count, junction count and gel coverage (Figure 3). Most importantly, the response was consistently non-monotonic: increasing nutrient concentration did not lead to a corresponding progressive increase in organised rhizomorph development.

**Figure 3.**
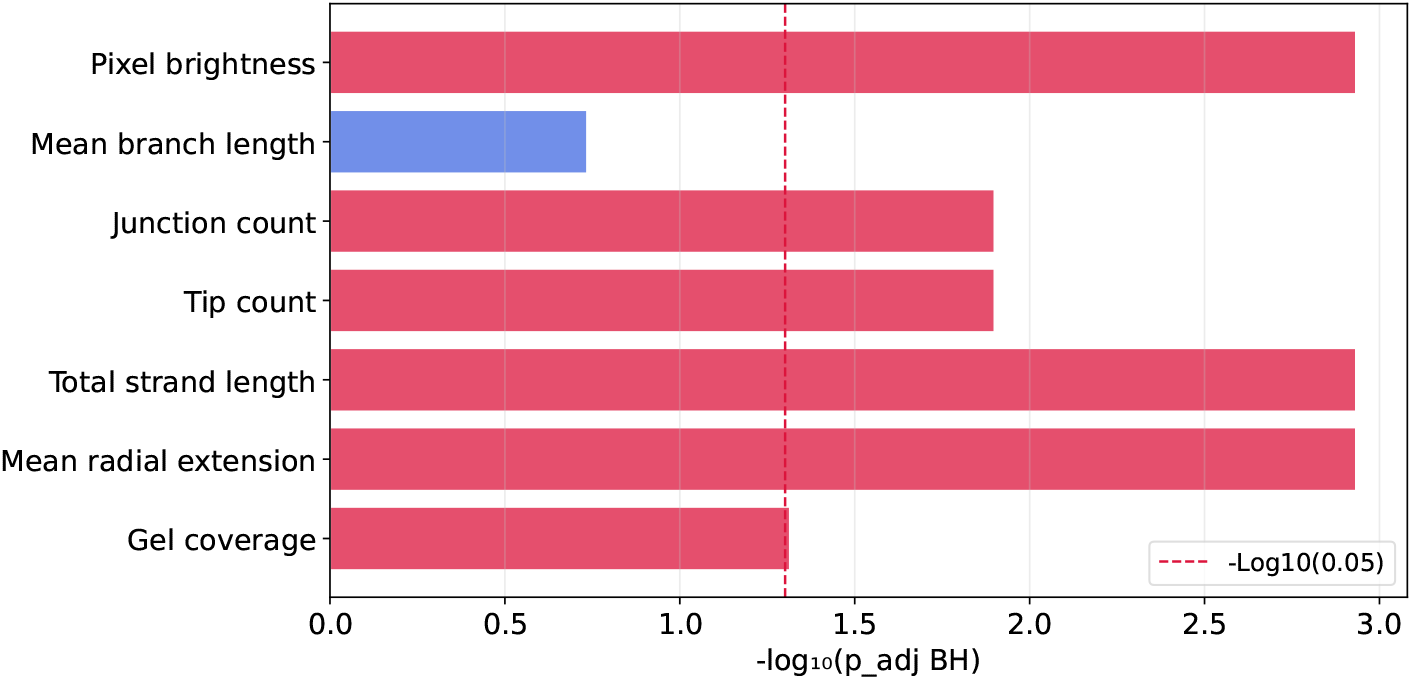
Statistical evidence for nutrient-dependent differences in fungal growth and network morphology. Permutation-based Kruskal–Wallis tests compared the four nutrient-density treatments at the final shared timepoint, with *p*-values adjusted across metrics using the Benjamini–Hochberg procedure; bar length represents *−* log_10_(*p*_adj_). Nutrient density significantly affected mean radial extension, total strand length, mean pixel brightness, tip count, junction count and gel coverage, demonstrating broad effects on both network extent and morphology.

#### Colonisation kinetics reveal a trade-off between rapid exploration and eventual coverage

Mean gel-coverage trajectories began to diverge after approximately 400 h (Figure 4a). The moderately nutrient-limited condition (level *−*1) expanded rapidly but plateaued at approximately 20% coverage, whereas the enriched conditions continued to increase later in the experiment. By approximately 900 h, level +1 had reached the greatest observed mean coverage, while level +2 remained intermediate. The most strongly nutrient-limited condition, level *−*2, showed the most restricted and irregular development.

**Figure 4.**
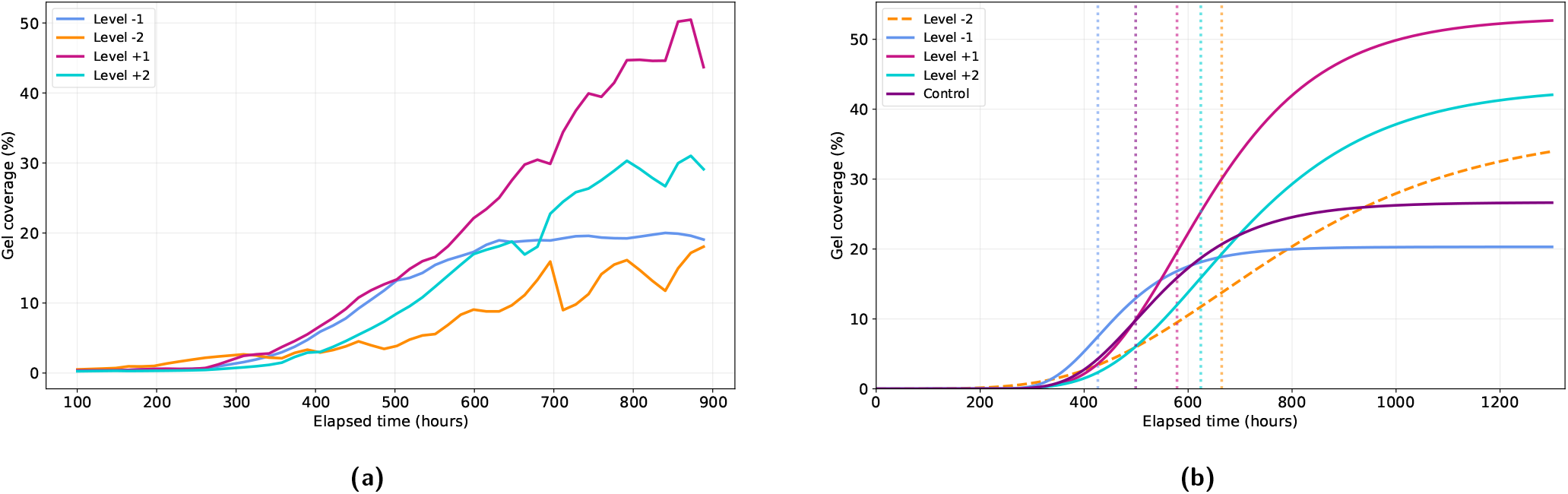
Nutrient-dependent colonisation dynamics. (a) Mean gel coverage over elapsed time across the four nutrient-density treatments, calculated as the fraction of the analysed dish (b) Mean Gompertz growth curves reconstructed from the arithmetic means of the retained replicate-level fitted parameters (*A, k* and *t*_*i*_); vertical dotted lines indicate the corresponding mean inflection times. Replicate fits were retained when *R*^2^ *≥* 0.70 and the projected asymptote satisfied *A ≤* 100%. The standard-nutrient control from the separate light-exposure experiment is shown for descriptive comparison only. Level *−* 1 showed the earliest inflection and highest growth-rate constant but the lowest fitted asymptote, whereas levels +1 and +2 developed later and reached higher fitted asymptotes. Estimates for level *−* 2 were highly variable and should therefore be interpreted cautiously.

Gel coverage differed significantly across the nutrient gradient in the omnibus Kruskal–Wallis analysis (*H* = 7.69, *p*_raw_ = 0.0415), and the effect remained significant after Benjamini–Hochberg multiple-testing correction (*p*_adj_ = 0.0484). The location of the apparent maximum at level +1 rather than at the highest nutrient concentration further suggests that the response was not simply dose dependent.

Gompertz modelling further distinguished the temporal growth strategies of the four conditions (Figure 4b; Table 1). Level *−* 1 showed the earliest mean inflection point, at approximately 426 h, the shortest lag phase, ending at approximately 331 h, and the highest fitted growth-rate constant, approximately 11.0 *×* 10^*−*3^ h^*−*1^. Despite this rapid early development, it reached the lowest fitted asymptote, approximately 20.3%. Levels +1 and +2 developed more slowly, with mean inflection points of approximately 579 and 624 h, respectively, but reached substantially higher fitted asymptotes. Level *−* 2 showed the slowest fitted dynamics, including the latest inflection point and lowest growth-rate constant. These parameters should, however, be treated cautiously because only four of five level *−* 2 samples yielded usable fits and those fits were highly variable (see Supplementary Figure 2a).

**Table 1.** Gompertz growth parameters for the four nutrient-density treatments and the standard-nutrient reference condition (mean *±* SD). The standard condition originates from the separate light-exposure experiment and is shown for descriptive comparison. Parameters were retained for fits with *R*^2^ *≥* 0.70 and projected asymptote *A ≤* 100%. *A* = asymptotic gel coverage (%); *k* = growth-rate constant (h^*−*1^); *t*_inflection_ = time of maximum growth rate (h); *t*_lag_ = lag-phase end (h) = *t*_inflection_ *−* 1*/k*. ^*†*^Only 4/5 samples yielded valid parameters (see Supplementary Figure 2a); interpret with caution. ^*‡*^One fit was excluded because its projected asymptote exceeded 100% (see Supplementary Figure 2b).

| Condition | $n$ fits / $n$ total | $A$ (%) | $k$ ($\times 10^{-3} \text{ h}^{-1}$) | $t_{\text{inflection}}$ (h) | $t_{\text{lag}}$ (h) | $R^2$ |
| --- | --- | --- | --- | --- | --- | --- |
| Control | 6 / 6 | $26.7 \pm 2.4$ | $8.28 \pm 1.44$ | $499 \pm 22$ | $376 \pm 11$ | 0.977 |
| Level $-2$ | 4 / 5<sup>†</sup> | $37.4 \pm 19.0$ | $3.67 \pm 0.75$ | $665 \pm 214$ | $384 \pm 171$ | 0.881 |
| Level $-1$ | 5 / 5 | $20.3 \pm 3.4$ | $10.99 \pm 2.85$ | $426 \pm 22$ | $331 \pm 11$ | 0.983 |
| Level $+1$ | 4 / 5<sup>‡</sup> | $53.2 \pm 11.1$ | $6.52 \pm 2.26$ | $579 \pm 30$ | $412 \pm 75$ | 0.911 |
| Level $+2$ | 5 / 5 | $43.2 \pm 18.7$ | $5.40 \pm 2.60$ | $624 \pm 120$ | $409 \pm 70$ | 0.957 |

The combined trajectories therefore suggest a nutrient-dependent trade-off between rapid exploration and eventual colonisation. Moderate nutrient limitation promoted earlier and faster expansion but a lower final coverage, whereas enriched conditions developed more slowly but ultimately colonised a larger fraction of the substrate. This interpretation is consistent with the ecological role of rhizomorphs as resource-translocating structures associated with fungal foraging [2, 6, 13, 17, 20]. Earlier studies have shown that rhizomorph initiation depends on nutrient status and that both nutrient deficiency and excess can restrict formation [16, 34]. The present results extend this picture by suggesting that nutrient availability also affects the temporal strategy of network development after initiation.

The high fitted asymptotes under nutrient-rich conditions should not, however, be interpreted as equivalent to increased production of organised rhizomorphs. Gel coverage quantifies all segmented fungal material irrespective of growth form. The accompanying structural and optical measurements show that high overall coverage could occur without a proportional increase in organised rhizomorphic network formation.

#### Network extent and spatial expansion peak at an intermediate nutrient level

Total strand length, mean radial extension and junction count provided complementary measures of organised network development and showed the same ordering at the final timepoint: level +1 *> −*1 *>* +2 *> −*2.

Level +1 produced the greatest mean total strand length, 974.5 *±* 544.8 mm, followed by level *−*1 at 681.1 *±* 114.4 mm, level +2 at 469.2 *±* 192.0 mm and level *−*2 at 178.9 *±* 126.0 mm (Figure 5a). Mean radial extension showed the same ordering, reaching 24.64 *±* 1.32 mm at level +1, 20.25 *±* 1.17 mm at level *−*1, 18.53 *±* 4.73 mm at level +2 and 14.73 *±* 3.03 mm at level *−*2 (Figure 5b). Level *−*1 generally led during the earlier stages of development, whereas level +1 overtook it during later growth. Thus, the condition producing the fastest early expansion was not the same as that producing the greatest eventual network extent.

**Figure 5.**
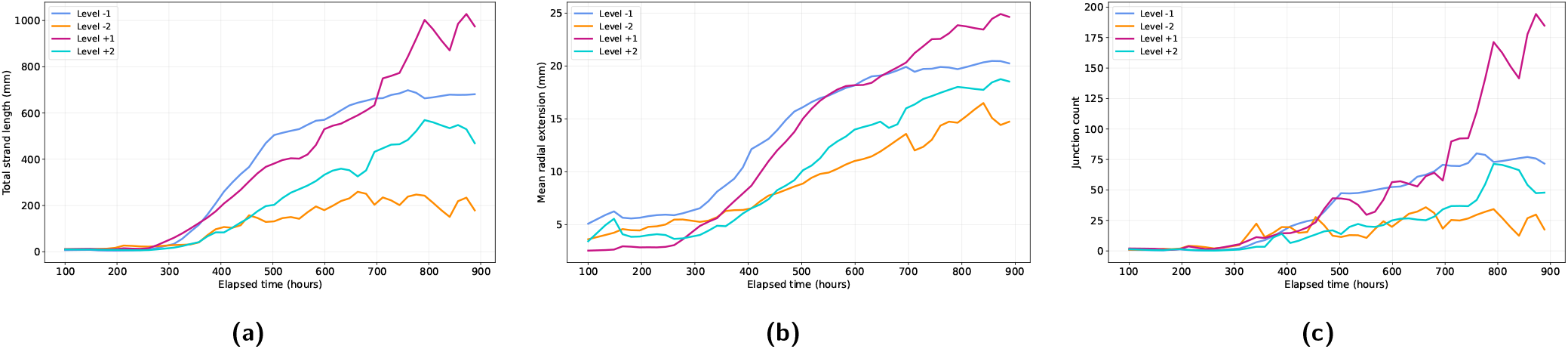
Nutrient-dependent development of organised rhizomorph networks. Metrics were derived from the segmented fungal regions and their skeletonised network representations over time: (a) total strand length quantifies overall network extent, (b) mean radial extension quantifies spatial expansion from the inoculation centre, and (c) junction count provides a measure of projected branching complexity. Level *−*1 led during early network development, whereas level +1 ultimately produced the greatest network extent and spatial expansion; further enrichment to level +2 did not increase organised network development, demonstrating a non-monotonic nutrient response.

Both total strand length and mean radial extension varied significantly among nutrient levels after correction for multiple comparisons (*H* = 12.78, *p*_adj_ = .0012 and *H* = 13.83, *p*_adj_ = .0012, respectively). Pairwise comparisons showed particularly clear separation of the severely nutrient-limited condition from levels *−*1 and +1, while level +1 also showed greater radial extension than both level *−*1 and level +2.

Junction count followed the same overall ranking (Figure 5c), with the highest mean observed at level +1. Its interpretation is less straightforward, however, because variability at level +1 was exceptionally large and no individual pairwise comparison remained significant after correction despite a significant omnibus test (*H* = 10.42, *p*_adj_ = .0126). In addition, validation of the skeletonisation procedure showed substantially greater condition-dependent error for junction count than for total strand length. The exact magnitude of differences in branching complexity should therefore be interpreted more cautiously than the agreement among total strand length and radial extension.

Taken together, the structural measurements support a non-monotonic response to nutrient availability. Severe nutrient limitation strongly restricted organised network development. Moderate limitation promoted rapid and comparatively consistent exploratory growth. Intermediate enrichment ultimately supported the greatest spatial expansion and network extent, although with substantial replicate-to-replicate variability. Increasing nutrient concentration further did not enhance organised network development and instead reduced total strand length and radial expansion relative to level +1.

This pattern is consistent with the idea that rhizomorph formation reflects a balance between resource acquisition and local substrate exploitation. When nutrients are limited but sufficient to sustain growth, investment in long-distance exploratory structures may be advantageous. At higher local nutrient availability, the benefit of such exploration may decrease, while resources can instead support denser or less spatially organised fungal growth. The present experiment does not directly measure fungal resource allocation, but the structural and optical data are consistent with such a shift in growth strategy.

#### Optical appearance changes under nutrient-rich conditions

Mean pixel brightness also differed significantly among nutrient treatments and diverged strongly during the later stages of growth (Figure 6a). The lower-nutrient conditions remained comparatively dark throughout the experiment, whereas levels +1 and +2 became progressively brighter from approximately 300–400 h onwards. The final-timepoint images showed a corresponding visual transition towards more diffuse and less strongly pigmented fungal growth in the nutrient-rich conditions (Figure 6b).

**Figure 6.**
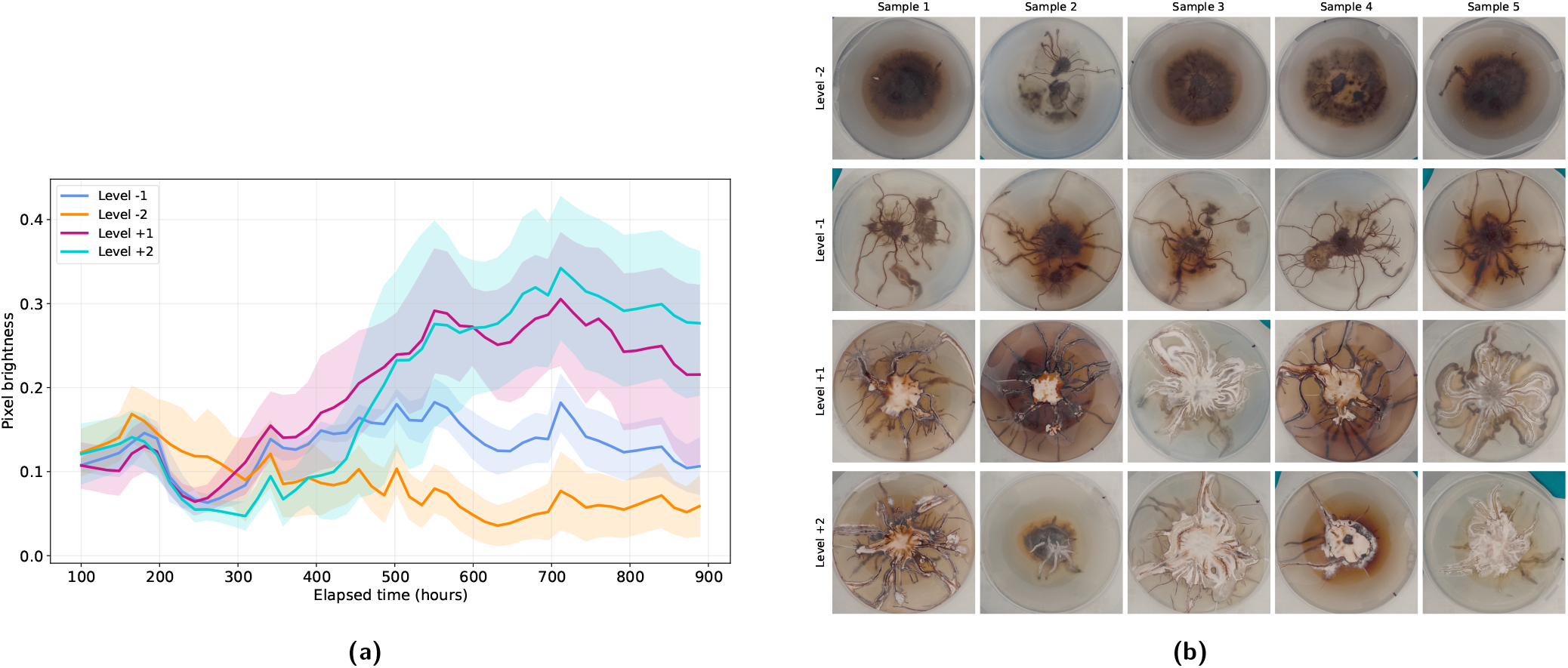
Nutrient-dependent changes in fungal optical appearance and growth morphology. (a) Mean brightness was calculated from normalised grayscale intensity within the segmented fungal region at each timepoint. (b) Final-timepoint plate images show the corresponding colony morphologies and were uniformly gamma-corrected (*γ* = 0.5) for visualisation. Higher nutrient conditions became progressively brighter and visually more diffuse during later growth, coinciding with high overall coverage but comparatively less organised network development. This pattern is consistent with an apparent shift in growth morphology, although brightness is an optical proxy and does not directly establish melanisation state or fungal structure identity.

The two nutrient-limited conditions should nevertheless not be treated as morphologically equivalent. Level *−*1 produced a clearly developed and coherent rhizomorphic network, whereas level *−*2 showed limited and frequently poorly defined growth. Its low brightness therefore cannot by itself be interpreted as evidence of a mature or highly organised rhizomorph phenotype. Rather, the combined structural and optical measurements indicate that level *−*2 restricted overall development, while level *−*1 produced the most rapid and consistent exploratory network.

At levels +1 and +2, increasing brightness coincided with the emergence of visually less-defined fungal structures and with a divergence between total coverage and organised network measurements. This pattern is consistent with an apparent transition towards less-pigmented or less-structured forms of mycelial growth under nutrient-rich conditions. Such a transition would provide a plausible explanation for why high nutrient availability increased total fungal coverage without producing a proportional increase in strand length, radial extension or branching organisation.

One possible biological interpretation is that abundant local resources reduce the need to invest in long-distance exploratory rhizomorphs, favouring more locally consolidating fungal growth. This interpretation remains tentative. Pixel brightness is an indirect optical measurement rather than a direct assay of melanin concentration, and it may also be influenced by strand thickness, imaging conditions and the underlying gel. Similarly, the image data cannot independently establish the anatomical identity of the brighter structures. The results therefore support a nutrient-dependent change in optical appearance and apparent growth morphology, but do not by themselves demonstrate altered melanisation or differentiation.

Overall, the nutrient experiment indicates that fungal coverage and organised rhizomorph development are related but distinct aspects of growth. Intermediate enrichment produced the greatest eventual network extent, whereas the highest nutrient concentration produced substantial fungal coverage without a corresponding increase in organised structural elaboration. Assessing multiple complementary metrics was therefore essential: coverage alone would have obscured the difference between total colonisation and the formation of an organised rhizomorphic network.

### 2.3. Exclusion of ambient light produces limited detectable changes in rhizomorph development

In contrast to the broad nutrient response, exclusion of ambient light produced no differences that remained statistically significant after correction for multiple comparisons (Figure 7). The illuminated and dark-grown samples showed broadly similar colonisation kinetics, optical development and timing of network formation. The clearest indications of a possible treatment response appeared in late-stage network architecture, but these differences remained descriptive rather than statistically conclusive.

**Figure 7.**
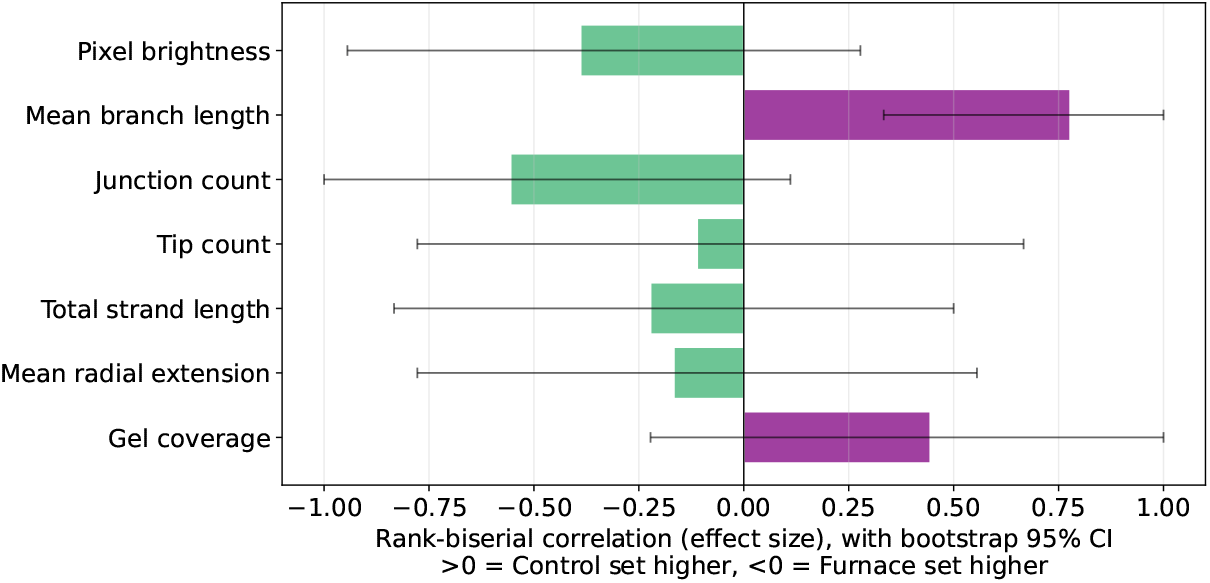
Effect of ambient-light exclusion on fungal growth and network morphology. Rank-biserial correlation coefficients were calculated from exact Mann–Whitney *U* tests comparing ambient-light and dark-grown samples at the final shared timepoint; error bars show bootstrap 95% confidence intervals, and significance was assessed after Benjamini–Hochberg correction across metrics. No measured metric differed significantly between conditions after correction, indicating that exclusion of ambient light produced comparatively limited detectable effects, with the clearest tendencies occurring in strand-level morphology.

#### Colonisation kinetics are largely conserved between illuminated and dark conditions

The two groups followed similar sigmoidal growth trajectories and developed in parallel until approximately 500 h (Figure 8a). During later growth, the dark-grown samples continued to expand somewhat further, plateauing at approximately 29–30% coverage compared with approximately 25–26% in the ambient-light condition. This corresponded to a moderate effect-size estimate but was not statistically significant after correction (*r* = .44, 95% CI [*−*.22, 1.00], *U* = 10, *p*_raw_ = .240, *p*_adj_ = .542).

**Figure 8.**
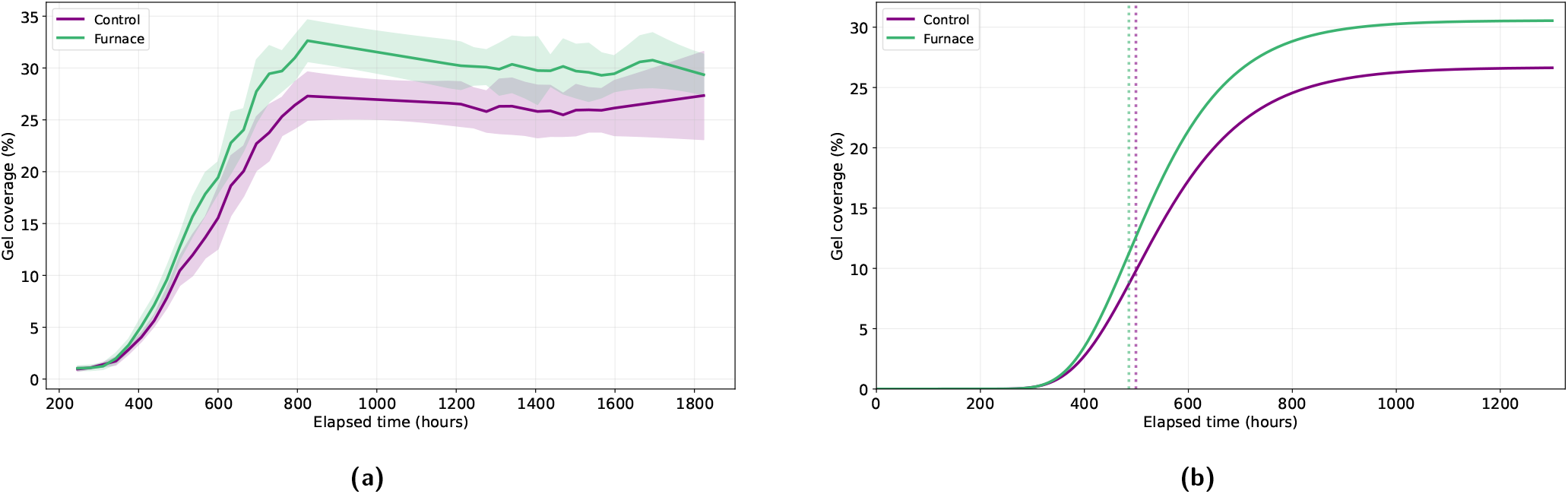
Colonisation dynamics under ambient-light and dark-grown conditions. (a) Mean gel coverage over elapsed time, calculated from the segmented fungal area of each dish. (b) Mean Gompertz growth curves reconstructed from the arithmetic means of the retained replicate-level fitted parameters; vertical dotted lines indicate the corresponding mean inflection times. The two conditions showed closely matched growth timing, while dark-grown samples reached a somewhat higher fitted asymptotic coverage. The observed coverage difference was not statistically significant after multiple-testing correction.

Gompertz modelling similarly showed closely matched growth dynamics (Table 2, Figure 8b). The dark-grown condition reached a somewhat higher fitted asymptote, 30.6 *±* 2.0% compared with 26.7 *±* 2.4%, and had a marginally higher fitted growth-rate constant (9.04 vs. 8.28 *×* 10^*−*3^ h^*−*1^). Its mean inflection point occurred slightly earlier, at approximately 486 h compared with 499 h. In contrast, lag phases were almost identical, approximately 373 and 376 h, indicating that removal of ambient light did not substantially alter the initiation or early acceleration of colonisation.

**Table 2.**
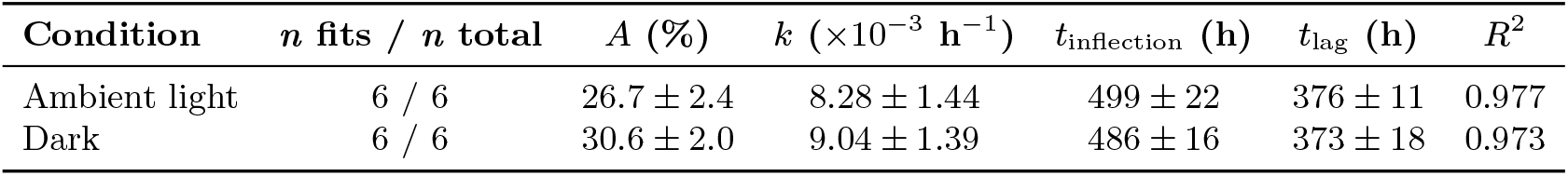
Gompertz growth parameters for ambient-light and dark-grown conditions (mean *±* SD). Parameters were retained for fits with *R*^2^ *≥* 0.70 and projected asymptote *A ≤* 100%. *A* = asymptotic gel coverage (%); *k* = growth-rate constant (h^*−*1^); *t*_inflection_ = time of maximum growth rate (h); and *t*_lag_ = lag-phase end (h) = *t*_inflection_ *−* 1*/k*.

These results provide little evidence that exclusion of ambient light fundamentally changed the broader colonisation programme under the conditions examined. This contrasts with earlier reports that darkness promotes rhizomorph formation and that illumination suppresses elongation in *Armillaria* [9, 37]. Several differences may account for this discrepancy, including fungal species, cultivation medium, developmental stage and the illumination regime itself.

In particular, the illuminated samples in the present experiment were exposed to ambient laboratory daylight rather than to a defined light intensity and photoperiod. The experiment should therefore be interpreted as a comparison between ambient-light and dark storage conditions rather than as a controlled light-dose experiment. The findings do not demonstrate that light generally has no effect on rhizomorph development; instead, they indicate that removing the ambient light exposure used here did not produce a large or statistically detectable alteration of colonisation kinetics.

#### Optical development is similar under both light environments

The optical trajectories of the two groups were also closely aligned (Figure 9). Mean brightness increased during network expansion, reached a shared peak of approximately 0.20 at around 800 h, and subsequently declined towards approximately 0.08–0.10 as growth approached its plateau. Standard-deviation bands overlapped through most of the observation period.

**Figure 9.**
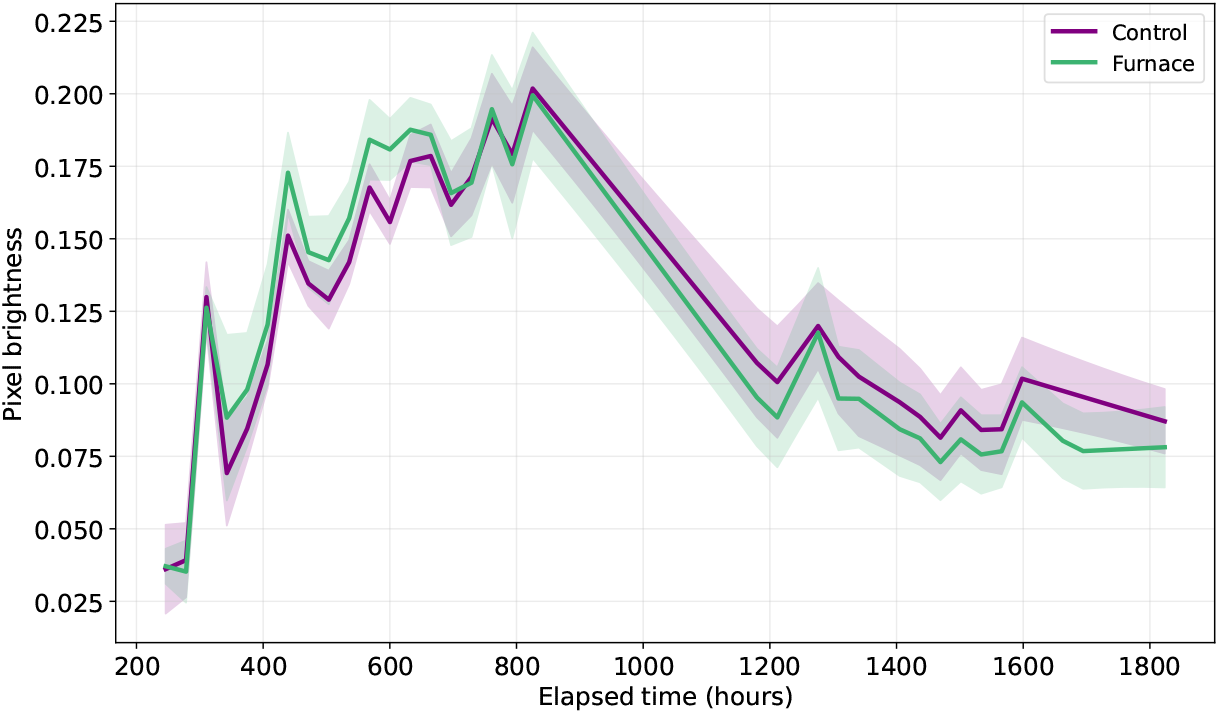
Optical development under ambient-light and dark-grown conditions. Mean brightness was calculated from normalised grayscale intensity within the segmented fungal region at each timepoint. Both conditions followed nearly overlapping trajectories, with increasing brightness during active expansion followed by progressive darkening as growth approached its plateau; the small late-stage divergence was not statistically significant, providing little evidence for a pronounced light-dependent optical response.

A small late-stage divergence was observed, with dark-grown samples reaching a slightly lower mean brightness than illuminated controls, but this difference was not statistically significant after correction (*r* = *−*.39, 95% CI [*−*.94, .28], *U* = 25, *p*_raw_ = .310, *p*_adj_ = .542). The broad similarity of the trajectories therefore provides little evidence for a pronounced light-dependent change in optical development.

The shared increase and subsequent decrease in brightness may instead represent a general feature of colony development. Newly extending regions appear optically lighter during active expansion, followed by progressive darkening as growth slows. Because brightness was not measured as a direct biochemical marker, this pattern cannot be attributed exclusively to melanisation. Nevertheless, the close agreement between treatments indicates that the temporal optical development of the colonies was largely preserved after exclusion of ambient light.

#### Possible differences are concentrated in late-stage network architecture

The strongest indications of a possible response to the light environment occurred in branch-level morphology during the later stages of network development (Figures 10a and 10b). At the final timepoint, dark-grown networks retained longer mean branches, approximately 5.51 *±* 0.77 mm compared with 4.49 *±* 0.46 mm under ambient light. The corresponding uncorrected comparison was significant (*U* = 4.0, *r*_rb_ = 0.78, *p*_raw_ = .026), but the difference did not remain significant after Benjamini–Hochberg correction (*p*_adj_ = .182).

**Figure 10.**
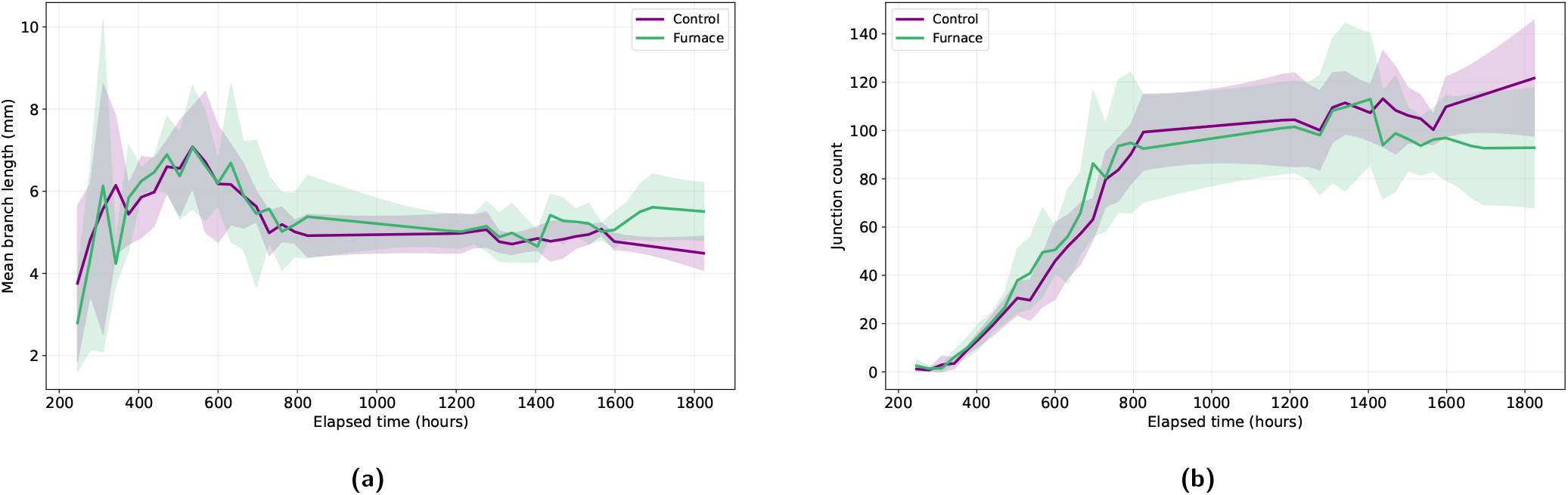
Network architecture under ambient-light and dark-grown conditions. Skeleton-derived trajectories show (a) mean branch length and **(b)** junction count as complementary measures of projected network subdivision and branching. During later development, dark-grown networks tended to retain longer branches and fewer junctions, consistent with a somewhat coarser network architecture. Neither difference remained significant after multiple-testing correction, so the pattern should be interpreted as a descriptive tendency rather than a demonstrated treatment effect.

Junction count showed the opposite tendency. Ambient-light networks contained approximately 121.7 *±* 26.3 junctions at the final timepoint, compared with 92.8 *±* 27.3 under dark conditions. This difference was also not statistically significant after correction (*U* = 28.0, *r*_rb_ = *−*0.56, *p*_adj_ = .462). The two observations are nevertheless internally consistent: a network containing fewer junctions will, all else being equal, be divided into fewer and therefore longer segments.

The dark-grown condition may consequently have produced a somewhat coarser late-stage architecture, characterised by longer uninterrupted branches and less subdivision, while the ambient-light condition tended towards a finer network. This interpretation should remain descriptive. Neither metric survived correction for multiple comparisons, the confidence interval for junction count crossed zero, and junction recovery is particularly sensitive to skeletonisation errors. The data therefore suggest a possible architectural tendency rather than a demonstrated treatment effect.

Directional growth was non-uniform in both illumination conditions, indicating that the asymmetry observed in individual colonies could not be attributed solely to the direction of ambient illumination. Because this analysis was exploratory and does not materially change the principal conclusion of the light experiment, it is more appropriately treated as a supplementary result.

## 3. Conclusions and Outlook

The principal outcome of this work is that, within the respective experimental designs, nutrient density produced substantially broader changes in rhizomorph network formation than the light environment, and that its influence was distinctly non-monotonic. Nutrient availability reshaped the timing, extent, spatial organisation and apparent morphology of growth, whereas exclusion of ambient light left the broader colonisation programme largely unchanged. The nutrient response is best understood as a trade-off among three competing outcomes: rapid exploration, organised network elaboration, and the production of alternative, less-structured fungal growth forms.

### Nutrient density and the exploration–elaboration trade-off

Moderate nutrient limitation (level *−*1) favoured exploration: it combined the earliest inflection, shortest lag and highest growth-rate constant with a comparatively low final coverage, yet still produced an extensive, radially expanded and reproducible network – a pattern consistent with rapid exploratory foraging under scarcity. Intermediate enrichment (level +1) favoured elaboration, supporting the greatest eventual strand length, radial extension and branching complexity, albeit with pronounced replicate-to-replicate variability. Crucially, further enrichment did not extend this trend: level +2 retained high coverage but formed a less elaborate network than level +1, while the lowest level ( *−*2) restricted development across all metrics. Among the nutrient levels tested, organised network elaboration was therefore greatest at intermediate enrichment rather than at the highest nutrient level.

The divergence between total coverage and organised structure at the nutrient-rich extreme is central to interpreting this trade-off. Levels +1 and +2 became progressively brighter and more variable during later growth, consistent with an apparent shift away from dark, well-defined rhizomorphic growth towards less-pigmented and less-structured fungal growth. Brightness alone cannot confirm the identity or melanisation state of these structures, but taken together with the coverage and network metrics it suggests that the additional coverage gained at high nutrient availability did not represent additional organised rhizomorph production. A plausible interpretation is that, once nutrients are abundant, the benefit of producing long-distance exploratory rhizomorphs may diminish and the organism may favour alternative, more consolidating growth forms – although confirming this requires methods able to distinguish structure types directly (see Limitations).

Methodologically, the combination of region-based, network-based and optical measurements was important because no individual metric fully captured the distinction between total fungal colonisation and organised rhizomorph development. In particular, the divergence between coverage and network metrics at higher nutrient levels illustrates why complementary measurements are required to distinguish overall fungal growth from the elaboration of an organised rhizomorphic network.

### Light exposure and the departure from prior reports

In contrast to the strong nutrient response – and contrary to earlier reports that darkness promotes rhizomorph development and that light inhibits growth in *Armillaria* [9, 37] – the light-exposure experiment provided little evidence that exclusion of ambient light altered the broader colonisation programme. Control and furnace samples shared similar lag and inflection times, closely overlapping brightness trajectories and consistently low inter-sample variability; the furnace condition reached only a marginally higher asymptotic coverage, and the clearest indications of a treatment response occurred in late-stage strand morphology, where furnace networks tended towards longer branches and fewer junctions, consistent with a comparatively coarser architecture. Furthermore, the similar progressive darkening observed in both groups is more consistent with a shared developmental trajectory than with a pronounced treatment-specific optical response.

This departure from the literature may reflect differences in species, cultivation conditions, light intensity, exposure duration, or the growth characteristics quantified across studies. Rather than a null result to be set aside, it represents a specific, testable discrepancy that warrants targeted re-examination under stringently controlled illumination. Consistent with a limited role for the light environment under the conditions tested, directional growth was nonuniform under both illuminated and dark conditions, indicating that the observed directional asymmetry was not driven solely by the direction of illumination but may instead reflect stochastic variation or local substrate heterogeneity.

### Limitations

Several limitations bound these conclusions. The sample size was small (*n* = 5–6 per condition), which limited statistical power, widened several effect-size confidence intervals, and left some apparent differences – including the higher furnace coverage and several nutrient contrasts – short of significance after correction for multiple comparisons. Inter-replicate variability was particularly high at more extreme nutrient levels (levels *−*2, +1 and +2), where growth departed most strongly from a well-defined rhizomorph network.

The optical and topological metrics are also indirect. Pixel brightness is a proxy for optical pigmentation rather than a direct measurement of melanin content, and the segmentation pipeline quantifies coverage and skeletal structure without classifying organised rhizomorphs and diffuse fungal growth as distinct biological categories. The inferred morphological transition at higher nutrient densities therefore remains an image-based interpretation. Two-dimensional skeletonisation cannot distinguish true fusion from strand contact or overlap, making junction- and tip-derived metrics less reliable than coverage, radial extension and total strand length. Conclusions based on the exact magnitude of branching complexity should therefore be interpreted accordingly.

Gompertz-derived parameters should likewise be interpreted cautiously for irregular growth, particularly at level *−*2, where fits were highly variable. The model provides a useful common description of sigmoidal colonisation dynamics but is less informative where growth departs substantially from this form.

Finally, the light and nutrient experiments differed in duration and experimental design. Gompertz fits permit descriptive comparison across them, but their raw effect magnitudes cannot be compared directly. Conclusions regarding the broader influence of nutrient density relative to the light environment therefore refer to the pattern of responses observed within the respective experimental designs rather than to a direct factorial comparison.

### Outlook

The most critical next step follows directly from the study’s original motivation. This work characterised how cultivation conditions shape rhizomorph network formation but did not measure mechanical performance; establishing whether the growth strategies identified here – particularly the well-organised networks formed under moderate and intermediate nutrient conditions – translate into a measurable mechanical advantage is therefore the necessary next step towards using rhizomorphs as reinforcement in mycelium-based composites.

The framework is well suited to that goal. Replacing manual capture with automated imaging would enable larger sample sizes, finer temporal resolution and the systematic screening of further variables such as temperature and geometric confinement. Future nutrient experiments should additionally include a contemporaneous standard-nutrient control and greater resolution across the range in which the strongest organised network development was observed. Likewise, the light experiment should be repeated under defined illumination intensity, spectrum and photoperiod to determine whether the discrepancy with previous studies persists under more stringently controlled conditions. A reduced and validated set of high-confidence metrics, complemented by methods capable of distinguishing organised rhizomorphs from alternative fungal growth forms, would further improve the robustness and biological interpretability of the framework.

Looking further ahead, linking cultivation conditions, network morphology and mechanical properties within a common quantitative framework could enable data-driven optimisation of rhizomorph architectures for application-specific material properties. Establishing such structure–property relationships would provide a stronger basis for assessing whether engineered rhizomorphic networks can improve the mechanical performance of mycelium-based composites and, ultimately, contribute to the development of higher-performance bio-derived materials. Other directions can include the use of, or training of, artificial intelligence networks [4, 35]. For instance, deeper insights into the complex behavior of biological systems can help us improve or inform the development of massively scaled agentic AI systems, for instance, or provide input into new learning algorithms.

## 4. Materials and Methods

### Fungal material and cultivation

*Armillaria gallica* rhizomorphs were obtained from the US Forest Service research centre *(Madison, USA)* and cloned in the laboratory prior to the experiments. Cultures were grown on potato dextrose agar (PDA; *Becton, Dickinson and Company, USA*) in distilled water *(ChemWorld, USA)*, prepared per the manufacturer’s recipe. Aniline blue diammonium salt *(Sigma-Aldrich, USA)* was added to enhance rhizomorph contrast against the gel and improve segmentation. For the nutrient-density experiment, four additional media were prepared with PDA concentrations scaled by factors of 0.25, 0.5, 1.5 and 2 relative to the standard (level 0); additional agar powder *(Thermo Fisher Scientific, USA)* maintained gel stability at reduced PDA (Table 3). Media were heated to 60 °C under continuous mixing until dissolved and sterilised in a pressure cooker.

**Table 3.** Composition of PDA media used to generate the nutrient-density treatments. Factors denote PDA concentration relative to the standard (level 0; factor 1) medium. Additional agar was added to the reduced-PDA media to maintain gel stability.

|  | *Level -2* | *Level -1* | *Level 0* | *Level +1* | *Level +2* |
| --- | --- | --- | --- | --- | --- |
| *Factor* | *0.25* | *0.5* | *1* | *1.5* | *2* |
| Water | 250 g | 250 g | 250 g | 250 g | 250 g |
| Potato dextrose agar | 3.0 g | 6.1 g | 12.2 g | 18.3 g | 24.4 g |
| Agar powder | 3.5 g | 2.3 g | — | — | — |
| Aniline blue | 0.063 g | 0.063 g | 0.063 g | 0.063 g | 0.063 g |

Set gel was reheated, rested to limit condensation, and 20 ml dispensed into Petri dishes (100 *×* 20 *mm, Corning, USA;* 100 *×* 15 *mm, VWR, USA)* in a sterile flow hood. A ~ 10 mm^2^ piece of a previously cultivated rhizomorph, cut with a sterile single-use scalpel, was placed centrally, and dishes were sealed at the outer edge with parafilm to maximise the observable area. Cultivation was at room temperature (20 °C) and 37% humidity. Light-exposure “furnace” samples were sealed from light in a furnace held at the same temperature as the illuminated set; all illuminated samples were grown under ambient daylight without controlled illumination (Table 4).

**Table 4.**
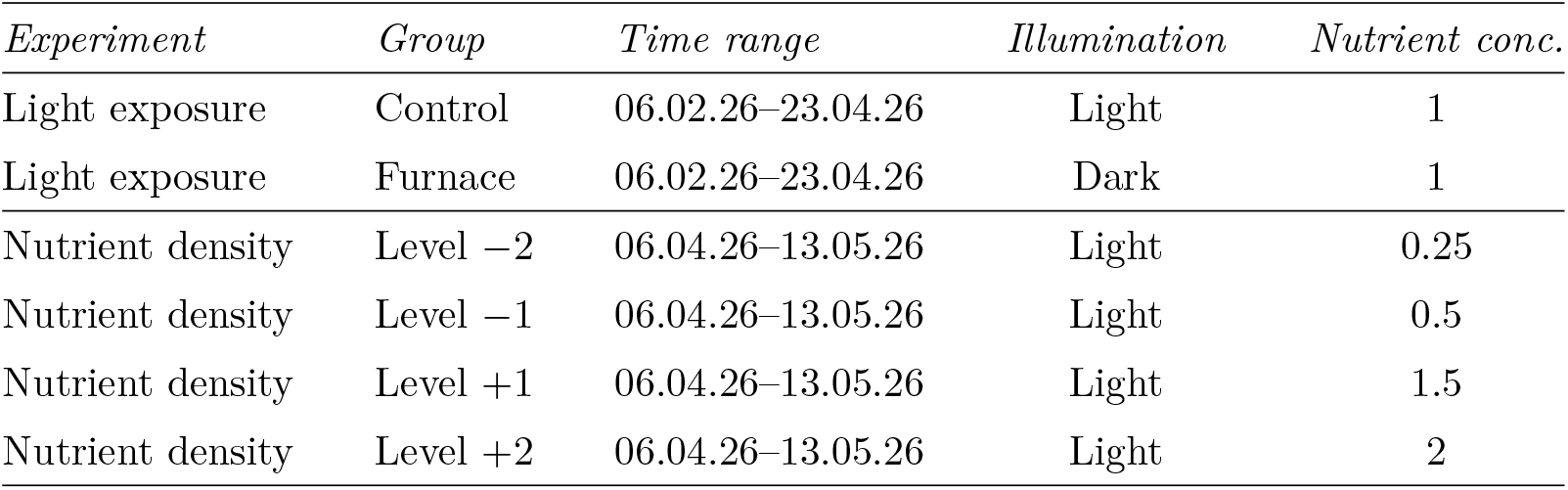
Experimental design and cultivation conditions for the light-environment and nutrient-density experiments. Nutrient concentration is expressed relative to the standard PDA concentration (factor 1). The two experiments were conducted separately and over different observation periods.

### Imaging

Samples were imaged manually in a lightbox *(Takerers, China)* using an iPhone 15 *(Apple, USA)* via Adobe Lightroom *(Adobe, USA)*, at ISO 40, shutter speed 1*/*200 s, 37 cm distance and 2*×* optical zoom. Imaging frequency ranged from daily early in each experiment to weekly in the later phase.

### Image segmentation

Segmentation used the Segment Anything Model 3 *(SAM 3; Meta AI)* [7] applied directly via the text prompt “dark rhizomorphs” without task-specific training. Each image was first preprocessed to improve strand visibility: the dish was localised, contrast enhanced by contrast-limited adaptive histogram equalisation and unsharp masking tuned to the gel type, and uniform background suppressed by a relative-darkness filter. The enhanced image was cropped to the dish, upscaled and passed to SAM 3; the highest-confidence masks (score *>* 0.5) were combined and clipped to the dish boundary to form the final binary mask. Masks that failed segmentation – no rhizomorph detected, or the entire background classified as such, occurring predominantly at early timepoints – were manually corrected (Supplementary Table 1).

Segmentation was validated against manually drawn ground-truth masks *(napari)* for 20 randomly selected images per condition, using pixel-level metrics for all masks and skeleton-based metrics for genuine networks only ( *≥* 10 ground-truth branches). Region overlap was reliable across conditions (median Dice 0.758–0.875, rising with nutrient level), with a mild over-segmentation tendency (precision below recall). Topological recovery was more variable and did not track pixel accuracy: on standard conditions tip, branch, junction and total-length counts deviated by *≤* 7% from ground truth, but under nutrient extremes junction- and branch-count biases were large and direction-dependent (e.g. junctions +60% at level *−*2, +41% at level +1, *−*16% at level *−*1). The pipeline is therefore suitable for area-based quantification, while topological metrics – particularly junction and tip counts at levels *−*2 and +1 – should be interpreted with corresponding caution (Supplementary Table 2).

### Growth quantification and network metrics

All analyses were performed in Python with scikit-image and OpenCV. In each image the dish was localised by a circular Hough transform and its known diameter (90 mm) used to derive a pixel-to-millimetre scale *s* = 90*/*(2 *r*_px_ *·*0.95); a group-wise median radius standardised the scale across frames of a sample, and all measurements were restricted to the inner 95% of the dish to exclude edge artefacts.

From each mask, we quantified colony extent, spatial expansion, network topology and optical appearance. Gel coverage and radial extension were calculated directly from the segmented region, while network architecture was derived from the final one-pixel-wide skeleton. Total strand length, branch-length statistics, tip and junction counts were extracted from the skeleton, and strand thickness was estimated from the Euclidean distance transform sampled along it. Pixel brightness was calculated from the normalised grayscale intensity of the segmented fungal region and used as an optical proxy for pigmentation and melanisation. Directional growth was quantified from the distribution of segmented pixels across twelve equal 30° angular sectors. Timestamps parsed from image filenames were used to express all metrics as functions of elapsed time. The complete definitions, calculation procedures and interpretations of the metrics used in the analysis are summarised in Table 5.

**Table 5.** Image-derived metrics used to quantify fungal colonisation, rhizomorph network morphology and optical appearance. Symbols: *M* denotes the set of colonised pixels within the clipped dish; |*M*| its pixel count; (*c*_*x*_, *c*_*y*_) the dish centre; *r*_px_ the dish radius in pixels; *g*(*p*) *∈* [0, 1] the normalised grayscale intensity at pixel *p*; *S* the one-pixel-wide skeleton; *n*(*p*) the number of skeleton neighbours of pixel *p*; *s* the pixel-to-millimetre scale factor; and *D*(*p*) the Euclidean distance from skeleton pixel *p* to the nearest background pixel.

| Metric | Calculation procedure | Interpretation |
| --- | --- | --- |
| Mean pixel brightness<br>(normalised intensity, 0–1) | $\frac{1}{\vert M\vert } \sum_{p \in M} g(p)$ | Mean optical intensity of the segmented fungal region. Lower values indicate darker, more strongly pigmented growth, whereas higher values indicate less-pigmented growth and may also reflect alternative mycelial structures. |

Table 5 – *continued from previous page*
| Metric | Calculation procedure | Interpretation |
| --- | --- | --- |
| Mean branch length (mm) | $\frac{1}{k} \sum_{i=1}^k L_i$ | Mean length of skeleton branches between endpoints and junctions. Lower values indicate a more finely subdivided network, whereas higher values indicate longer uninterrupted branches. |
| Junction count (count) | Number of skeleton junctions identified from the final network | Number of branching points in the skeleton and therefore a measure of network branching complexity. |
| Tip count (count) | Number of terminal endpoints identified in the final skeleton | Number of free-growing strand ends; higher values indicate a greater number of active exploratory fronts. |
| Total strand length (mm) | $s \sum_{i=1}^k L_i$ | Summed length of all branches in the skeleton, representing the total extent of the organised fungal network. |
| Mean radial extension (mm) | $\frac{s}{\vert M\vert } \sum_{(x,y) \in M} \sqrt{(x - c_x)^2 + (y - c_y)^2}$ | Mean distance of colonised pixels from the inoculation centre, quantifying the spatial extent of outward growth. |
| Gel coverage (%) | $100 \cdot \frac{\vert M\vert }{\pi(0.95 r_{\text{px}})^2}$ | Percentage of the analysed dish area occupied by segmented fungal growth and the primary measure of overall colonisation. |
| Directional sector fraction (dimensionless) | $\frac{\vert \{(x,y) \in M : \theta_a \leq \theta < \theta_b\}\vert }{\vert M\vert }$, with twelve equal $30^\circ$ sectors | Fraction of total segmented growth occurring within a given angular sector. A spatially uniform colony has an expected fraction of approximately $1/12$ per sector; deviations indicate directional asymmetry. |

### Statistical analysis

Analyses used SciPy, statsmodels and scikit-learn, with *α* = 0.05 and a fixed random seed for all bootstrap and permutation procedures. Only timepoints shared by at least min(3, *n*) samples per group were included. Replicate numbers were *n* = 6 (control, furnace) and *n* = 5 (each nutrient level). All tests were two-sided.

Gel-coverage trajectories were fitted per replicate with a Gompertz model,

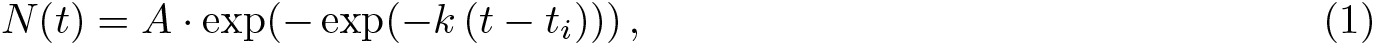

by bounded nonlinear least squares (trust-region reflective, 18-point multi-start; highest-*R*^2^ fit retained), where *A* is the asymptote, *k* the rate constant and *t*_*i*_ the inflection time; lag was *t*_lag_ = *t*_*i*_ *−* 1*/k*. Fits with *R*^2^ *<* 0.70 were excluded from condition-level summaries, and a Gompertz-over-linear *R*^2^ gain *>* 0.05 was taken to confirm non-linear growth; for the cross-level parameter tables, fits with a projected asymptote *>* 100% were additionally excluded as non-physical. Because the two experiments ran over different durations, only the fitted Gompertz curves – not raw final values – are compared across experiments; raw coverage plots retain each set’s original elapsed time.

Two-group comparisons (light exposure) used the exact Mann–Whitney *U* test at the final shared timepoint, with rank-biserial correlation *r* = 1 *−* 2*U/*(*n*_1_*n*_2_) and 95% bootstrap confidence intervals (2000 resamples). Multi-group comparisons (nutrient density) used Kruskal–Wallis tests with permutation-based *p*-values (1999 permutations). Exact two-sided Mann–Whitney comparisons were additionally calculated across all nutrient-level pairs for each metric to quantify pairwise differences, irrespective of omnibus significance. Kruskal–Wallis *p*-values were Benjamini–Hochberg adjusted across metrics, while pairwise *p*-values were adjusted across the six level comparisons within each metric. Directional uniformity (light-exposure experiment only) was assessed by a *χ*^2^ goodness-of-fit test against a uniform 12-sector expectation, per replicate and for the condition-level mean.

### Use of generative AI

During manuscript preparation, an LLM was used to assist with language editing and code editing for benchmark generation scripts. The authors reviewed and edited the output, verified the scientific content, and are responsible for the final manuscript.

## Supplementary Information

The Supplementary Information document provides additional details and methods.

## Author contributions

S.C.N.: Conceptualization, methodology, investigation, data curation, software, formal analysis, visualization, and writing - original draft. M.J.B.: Conceptualization, methodology, supervision, project administration, and writing - review and editing. Both authors contributed to interpretation of the results and approved the final manuscript.

## Code and data availability

The complete software environment and package versions used in the analyses are provided in Supplementary Table 3. All custom code used for image preprocessing, segmentation, growth quantification, statistical analysis and figure generation is publicly available in the project GitHub repository: https://github.com/lamm-mit/rhizomorphic-networks. All data is provided via https://huggingface.co/datasets/lamm-mit/rhizomorphic-networks-data.

