## Supplementary Information for "Nutrient Availability Regulates an Exploration-Elaboration Trade-off in Fungal Rhizomorph Networks"

This file contains Supplementary Note 1, Supplementary Figures 1–2 and Supplementary Tables 1–3. Items are referenced from the main text in the order in which they first appear.

### Supplementary Note 1: Gompertz fit retention

Gompertz parameters were retained only for replicates meeting two criteria: a fit quality of $R^2 \geq 0.70$, and – to avoid implausible extrapolation – a projected asymptote of $A \leq 100\%$. Under these criteria all six control replicates and all five replicates at levels $-1$ and $+2$ were retained. At level $+1$, one otherwise acceptable fit was excluded because its projected asymptote exceeded 100% (a clear artefact projecting $\approx 132\%$), leaving four of five. At level $-2$, only four of five replicates could be fitted at all: replicate 4 could not be fitted and was poorly described even by a linear model ($R^2_{\text{linear}} = 0.17$; Supplementary Figure S2a), and the four retained fits were of highly variable quality. Level $-2$ parameters are therefore reported as indicative rather than precise. Because the two experiments ran over different durations, only the fitted Gompertz curves – not raw final values – are compared across experiments.

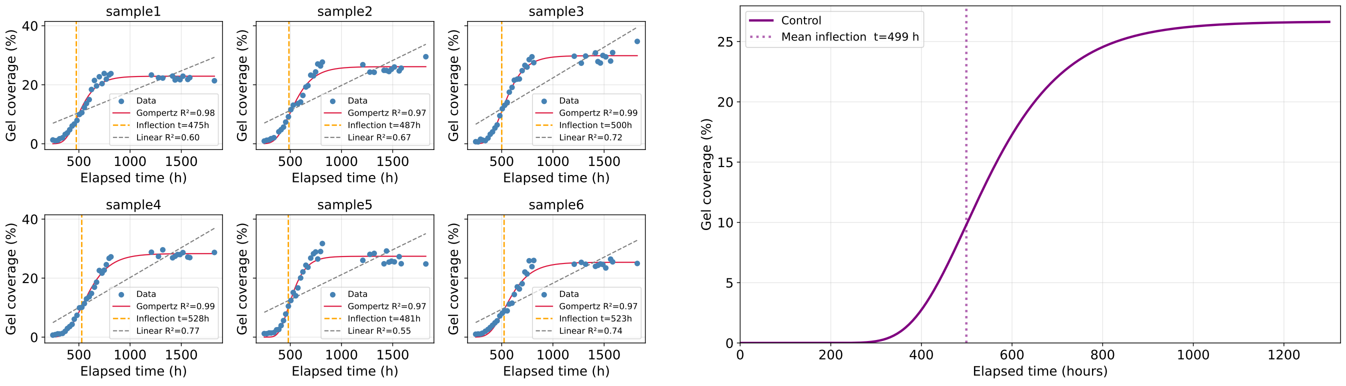


(a) (b)

**Figure S1: Gompertz modelling of gel-coverage dynamics under the control condition. (a)** Replicate-level Gompertz fits to observed gel-coverage trajectories. Blue points show measured gel coverage, red curves show the fitted Gompertz models, orange dashed lines indicate fitted inflection times, and grey dashed lines show the corresponding linear fits. All six control replicates were well described by the Gompertz model ($R^2 = 0.97$–$0.99$), with fitted inflection times ranging from approximately 475 to 528 h. **(b)** Mean Gompertz growth curve reconstructed from the arithmetic means of the retained replicate-level fitted parameters. The vertical dotted line indicates the mean fitted inflection time ($t_i \approx 499$ h). All six control fits satisfied the retention criteria of $R^2 \geq 0.70$ and projected asymptote $A \leq 100\%$.

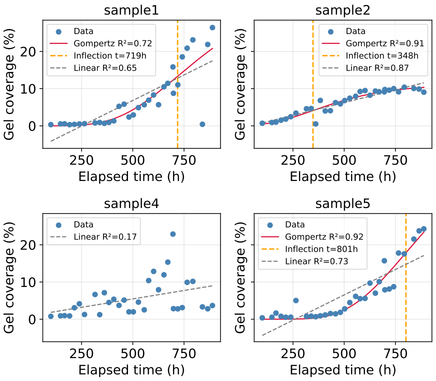


**(a)** Nutrient-density level $-2$.

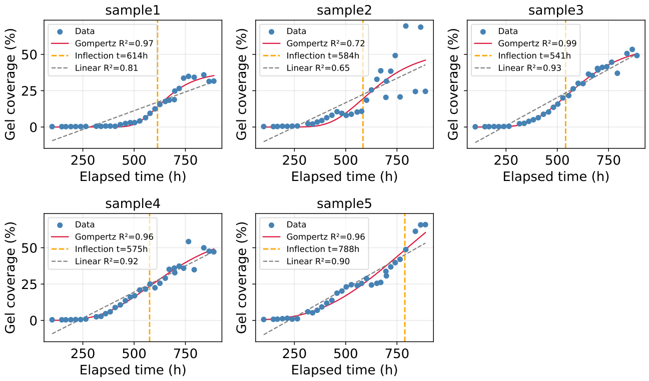


**(b)** Nutrient-density level $+1$.

**Figure S2: Per-replicate Gompertz fits to gel-coverage trajectories at nutrient-density levels $-2$ and $+1$.** Blue points show observed gel coverage (%) over elapsed time, red curves show the fitted Gompertz model $N(t) = A\exp(-\exp(-k(t-t_i)))$, orange dashed lines indicate fitted inflection times $t_i$, and grey dashed lines show the corresponding linear fits. Fits were obtained by bounded nonlinear least squares (see Methods). **(a)** At level $-2$, organised branching growth largely broke down, with colonies expanding as diffuse, darkening blobs. Only four of five replicates yielded usable fits; replicate 4 could not be fitted and was poorly described even by a linear model ($R^2_{\text{linear}} = 0.17$). The retained fits were of highly variable quality, so level $-2$ parameter estimates should be interpreted as indicative rather than precise. **(b)** At level $+1$, all five replicates achieved $R^2 \geq 0.70$, but one otherwise acceptable fit was excluded from parameter-level comparisons because its projected asymptote exceeded the retention criterion of $A \leq 100\%$, reaching approximately $132\%$. Consequently, four of five replicates were retained for parameter summaries; among these, Gompertz $R^2$ values ranged from $0.72$ to $0.99$, and fitted inflection times ranged from $541$ to $614$ h (mean $579$ h).

**Table S1: Dataset composition and mask-validation outcomes across experimental conditions.** "Valid masks" are segmentations accepted without intervention; "corrected masks" were manually post-processed, predominantly at early timepoints where SAM 3 either detected no rhizomorph or classified the entire background as foreground.

| | *Samples* | *Timepoints* | *Available images* | *Valid masks* | *Corrected masks* |
|---|---|---|---|---|---|
| Control | 6 | 36 | 216 | 204 | 12 |
| Furnace | 6 | 38 | 228 | 218 | 10 |
| Level $-2$ | 5 | 32 | 160 | 122 | 38 |
| Level $-1$ | 5 | 32 | 160 | 131 | 19 |
| Level $+1$ | 5 | 32 | 160 | 111 | 49 |
| Level $+2$ | 5 | 32 | 160 | 114 | 46 |

**Table S2: Segmentation performance against manually drawn ground truth.** Region-overlap, coverage and boundary metrics use all available matched masks ($n = 30$). Skeleton-structure metrics use genuine networks only (ground-truth final skeleton with $\geq 10$ branches; $n$ given per column) and are calculated from the final blob-aware, pruned skeletons. Segmentation values are based on raw, uncorrected SAM 3 output and therefore represent a conservative estimate of the quality of the production segmentations, in which isolated failures were manually corrected (Supplementary Table S1). Median coverage bias is the median per-image signed relative difference in predicted versus ground-truth area. For skeleton metrics, Δ% denotes the pooled signed deviation from ground truth, $100(\sum \text{pred}/\sum \text{GT} - 1)$.

| | Light exposure | | Nutrient density | | | |
|---|---|---|---|---|---|---|
| Metric | Control | Furnace | Level −2 | Level −1 | Level +1 | Level +2 |
| Dice | 0.849 | 0.887 | 0.749 | 0.781 | 0.857 | 0.748 |
| IoU | 0.743 | 0.802 | 0.640 | 0.681 | 0.773 | 0.668 |
| Precision | 0.816 | 0.883 | 0.711 | 0.740 | 0.853 | 0.775 |
| Recall | 0.898 | 0.901 | 0.902 | 0.882 | 0.907 | 0.837 |
| Specificity | 0.985 | 0.990 | 0.995 | 0.989 | 0.992 | 0.996 |
| Accuracy | 0.984 | 0.989 | 0.993 | 0.988 | 0.990 | 0.993 |
| Med. coverage bias (%) | +9.7 | +7.1 | +21.0 | +11.5 | +5.1 | −1.3 |
| $n$ (genuine) | 28 | 28 | 18 | 25 | 22 | 24 |
| Tips Δ% | −8.3 | −4.9 | +29.6 | −8.5 | +45.4 | +15.3 |
| Branches Δ% | −4.8 | −0.8 | +26.0 | −13.1 | +61.4 | +10.0 |
| Junctions Δ% | −3.8 | +0.5 | +25.3 | −14.4 | +66.3 | +7.8 |
| Total length Δ% | −2.8 | −6.1 | +8.6 | −9.0 | +9.7 | −10.8 |
| Mean branch len. Δ% | +1.5 | −5.0 | −3.6 | +4.1 | −28.7 | −6.8 |

**Table S3: Software and package versions used for the computational analyses.** A complete frozen environment specification, including all transitive dependencies, together with all analysis code, is available in the project repository (see Code Availability in the main text).

| *Software / package* | *Version* | *Purpose* |
|---|---|---|
| Python | 3.10.19 | Programming language |
| SAM 3 | 0.1.0 | Image segmentation model |
| Hugging Face Transformers | git `de306e8`[a] | SAM 3 implementation |
| PyTorch | 2.7.1 (CUDA 11.8) | Deep-learning backend |
| torchvision | 0.22.1 (CUDA 11.8) | Image transforms |
| OpenCV | 4.13.0.90 | Image preprocessing |
| scikit-image | 0.25.2 | Image processing |
| skan | 0.13.1 | Skeleton network analysis |
| napari | 0.6.6 | Manual ground-truth masks |
| NumPy | 1.26.4 | Numerical computing |
| SciPy | 1.15.3 | Curve fitting, statistical tests |
| statsmodels | 0.14.6 | Multiple-testing correction |
| scikit-learn | 1.7.2 | Statistical analysis |
| pandas | 2.3.3 | Data handling |
| Matplotlib | 3.10.8 | Visualisation |

[a] Installed from the development branch at the commit hash shown, which provided the SAM 3 implementation prior to a tagged release.
